# Efficient CRISPR/Cas9 mutagenesis for neurobehavioral screening in adult zebrafish

**DOI:** 10.1101/2021.02.01.429280

**Authors:** Dana Klatt Shaw, Mayssa H. Mokalled

## Abstract

Adult zebrafish are increasingly used to interrogate mechanisms of disease development and tissue regeneration. Yet, the prospect of large-scale genetics in adult zebrafish has traditionally faced a host of biological and technical challenges. Here, we describe an experimental pipeline that combines high-efficiency CRISPR/Cas9 mutagenesis with functional phenotypic screening to identify genes required for spinal cord repair in adult zebrafish. Using CRISPR/Cas9 dual-guide ribonucleic proteins, we show selective and combinatorial mutagenesis of 17 genes at 28 target sites with efficiencies exceeding 85% in adult F_0_ ‘crispants’. We find that capillary electrophoresis is a reliable method to measure indel frequencies, while avoiding the limitations of restriction enzyme-based genotyping. Using a quantifiable behavioral assay, we identify 7 single- or duplicate-gene crispants with reduced functional recovery after spinal cord injury. To rule out off-target effects, we generate germline mutations that recapitulate the crispant regeneration phenotypes. This study provides a platform that combines high-efficiency somatic mutagenesis with a functional phenotypic readout to perform medium- to large-scale genetic studies in adult zebrafish.

## INTRODUCTION

Zebrafish are a premier model to interrogate mechanisms of vertebrate biology. Embryonic and larval zebrafish are traditionally employed to probe vertebrate embryogenesis (Eisen 1996; Schier and Talbot 2005; Streisinger et al. 1981). But more recently, adult zebrafish are eminently used to model tissue physiology and disease mechanisms, including arthritis, scoliosis, cancer, blood disorders and undiagnosed diseases (Askary et al. 2016; Gray et al. 2020; Kaufman et al. 2016; Langenau et al. 2003; Van Gennip et al. 2018; Wangler et al. 2017). Due to their renowned regenerative capacity, adult zebrafish are widely used to uncover injury responses and repair mechanisms in multiple tissues such as fin, heart, and pancreas (Kroehne et al. 2011; Moss et al. 2009; Poss et al. 2002b; Tu and Johnson 2011; Vihtelic and Hyde 2000; Yurco and Cameron 2005). The remarkable capacity to regenerate neural tissues, including brain and spinal cord, is attracting a growing community of scientists into neurobehavioral studies in adult zebrafish (Becker and Becker 2015; Mokalled and Poss 2018; Orger and de Polavieja 2017). Thus, there is a pressing need to refine and expand the genetic and molecular toolkit for adult zebrafish research.

Developing zebrafish embryos are ideal for large-scale genetic screening. Zebrafish generate large clutches of transparent embryos, undergo rapid external development, and allow for direct visualization of phenotypes at embryonic and larval stages. These experimental advantages empowered forward genetic screening in nervous system development and behavior (Brockerhoff et al. 1995; Granato et al. 1996; Moens et al. 1996; Wolman et al. 2015). Conversely, studies of the much larger, non-transparent tissues of adult zebrafish have historically presented a number of technical and practical challenges. 1) Genetic screening requires the ability to grow large numbers of animals, preferably within a small footprint and at a relatively cheap cost. 2) Genetic screening involves multiple generations of breeding, with only a fraction of the resulting F_3_ animals displaying homozygosity. 3) The success of a genetic screen relies on obtaining highly penetrant and accessible phenotypic readouts. 4) Importantly, in the absence of conditional targeting approaches, genetic screening in adult zebrafish is restricted to genes that are dispensable for embryonic development. The limited ability to identify adult phenotypes, coupled with the spatial and technical demands of adult husbandry, have curtailed the use of adult zebrafish for high-throughput genetic studies.

A number of genetic screens have defied the challenges of large-scale genetics in adult zebrafish. These screens preferentially targeted accessible tissues such as skin pigmentation or fin regeneration, or sought easily discernable phenotypes in less accessible tissues such as sterility or scoliosis (Dosch et al. 2004; Gray et al. 2020; Haffter et al. 1996; Henke et al. 2017; Maderspacher and Nusslein-Volhard 2003; Wagner et al. 2004). Notably, temperature-sensitive genetic screens in adult zebrafish provided an additional advantage, by screening for mutations that bypass early development at permissive temperature but impair fin regeneration at non-permissive temperature (Johnson and Weston 1995; Oppedal and Goldsmith 2010; Poss et al. 2002a). While many of these adult genetic screens are non-saturating, they remain unfeasible in less accessible tissues such as the adult spinal cord, precluding large-scale genetic studies in adult zebrafish.

Over the past decade, increased accessibility to large-scale transcriptomics and the advent of CRISPR/Cas9 genome editing revolutionized reverse genetic studies in adult zebrafish. Standard mutagenesis protocols showed inconsistent efficiencies and variable phenotype penetrance, limiting their application to small-scale studies of single or double germline mutant lines (Burger et al. 2016; Gagnon et al. 2014; Hwang et al. 2013; Jao et al. 2013; Kotani et al. 2015; Shah et al. 2015; Varshney et al. 2015; Wu et al. 2018). Recent development of high-efficiency targeting protocols reinvigorated the prospect of larger scale, somatic mutagenesis in adult zebrafish (Hoshijima et al. 2019; Xu et al. 2020). Chemically synthesized CRISPR/Cas9 dual-guide ribonucleic protein (dgRNP) complexes were shown to reliably produce somatic mutations with >90% efficiency, and to mimic germline mutant phenotypes in transiently targeted, developing zebrafish embryos (Hoshijima et al. 2019). Yet, the efficiency and applicability of CRISPR/Cas9 dgRNP induced mutagenesis in adult zebrafish remains to be determined.

Here, we combined high-efficiency CRISPR/Cas9 mutagenesis with a functional neurobehavioral readout to identify genes necessary for spinal cord regeneration in adult zebrafish. A total of 17 genes, including 3 sets of duplicate paralogs and 1 pair of functionally redundant genes, were individually and combinatorially targeted. We found CRISPR/Cas9 dgRNPs eliminated more than 85% of wild-type gene copies in larval and adult crispants. Ten genes showed comparable mutagenesis rates between larvae and adults. For the 7 remaining genes, wild-type alleles were recovered at higher frequency in adult animals, suggesting a subset of mutant alleles are subject to negative selection in juvenile zebrafish. Using next generation sequencing, we found capillary electrophoresis effectively measures indel frequency in individual fish. Using swim capacity as a readout of motor function recovery, we identified 5 single genes, 1 gene duplicate pair, and 1 pair of functionally redundant genes that are required for functional spinal cord repair after spinal cord injury (SCI). Finally, we generated 5 germline mutations that recapitulated the histological and functional regeneration phenotypes of targeted crispants. Taken together, this study provides an experimental framework that combines high-efficiency somatic mutagenesis with a functional phenotypic readout to perform medium-to large-scale genetic studies in adult zebrafish.

## RESULTS

### Gene targeting by CRISPR/Cas9 dgRNPs

We used high-efficiency CRISPR/Cas9 mutagenesis to identify genes that direct spontaneous spinal cord repair in adult zebrafish (Hoshijima et al. 2019). Using previously generated transcriptomic datasets, we selected 14 genes that are upregulated in the spinal cord after injury (GEO accession # GSE164945) (Shaw et al. 2021, In Press). Genes of interest were filtered based on their biological function and previous characterization during embryonic development. For biological function, we preferentially targeted transcription factors that are enriched in regenerative glial cells after injury. To prevent early developmental lethality, we excluded genes that were previously associated with lethal phenotypes in embryonic or juvenile mutant animals (zfin.org). We also used the EMBL-EBI *Danio rerio* expression atlas to select genes that were either maternally supplied or not expressed at early zygotic stages (https://www.ebi.ac.uk/gxa/experiments/E-ERAD-475). To reduce the likelihood of genetic compensation between duplicate copies of the same gene (Postlethwait et al. 1998), gene duplicates were targeted alone and in tandem (Figure 1A). For each candidate gene, we designed 1 or 2 targeting dgRNAs that do not have any predicted off-target sites in the zebrafish genome (Labun et al. 2019). Overall, we targeted 17 genes at 28 target sites.

**Figure 1.**
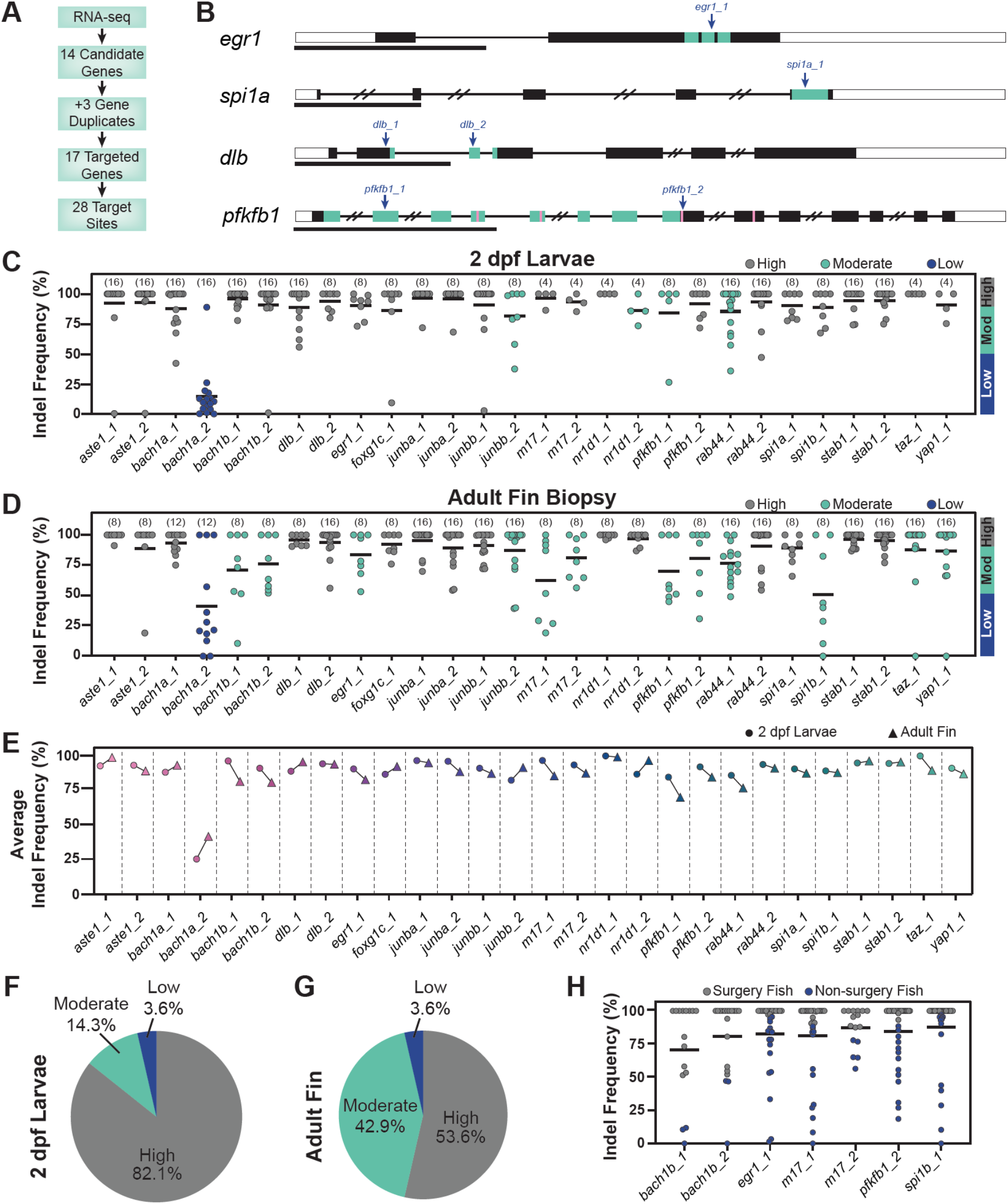
CRISPR/Cas9 dgRNPs effectively mutate candidate genes in adult zebrafish. **(A)** Schematic summary of gene targeting employed is this study. A total of 17 genes, including 3 pairs of gene paralogs, were targeted using 28 dgRNPs. **(B)** Representative schematics for dgRNP targeted genes. Shown are transcription factors *egr1* and *spi1a*, the *dlb* ligand, and the enzyme *pfkfb1*. Black boxes are exons, white boxes are UTRs, and lines are introns. Key domains are indicated in teal: zinc finger (*egr1*) and ETS (*spi1a*) DNA binding domains, EGF-like domain 1 (*dlb*), and 6PF2K domain (*pfkfb1*). The enzymatic active sites are indicated by pink lines in *pfkfb1*. dgRNP target sites are indicated by blue arrows. Scale bars represent 1 kilobase (Kb). **(C,D)** Targeting efficiency in dgRNP injected zebrafish crispants. Capillary electrophoresis was used to measure amplicon length following PCR based genotyping. Data points represent individual animals for each target site. Lines indicate means. Sample sizes are indicated between parentheses. 2 dpf larvae (C) or adult caudal fin (D) were analyzed. Indel frequency represents the percent of non-wild-type PCR amplicons relative to total PCR products amplified for each target site. Target sites are classified into 3 categories: high-efficiency (indel frequency >90%, gray), moderate-efficiency (indel frequency 50-90%, teal), and low-efficiency (indel frequency <50%, blue). **(E)** Comparison of dgRNP targeting efficiency between larval and adult (triangles) zebrafish. Data points represent average indel frequency for each target site. Circles and triangles indicate average indel frequency in 2dpf larva and 4 month adults, respectively. **(F,G)** Pie chart representation of dgRNP efficiency. Shown are the fractions of dgRNPs that fall into each category: high (gray), moderate (teal), and low (blue). Each pie chart represents 28 tested dgRNPs. **(H)** Capillary electrophoresis genotyping of adult animals in which indel frequency decreased between larvae and adults. Data points represent individual animals. Adult fish with indel frequency >90% (gray) were subjected to SCI and subsequent phenotyping. Because only 16 *taz_1/yap_1* dual-targeted animals survived to adulthood, all were genotyped in D.

To maximize the effect of small indels, we selected target sites within key domains (uniport.org), such as DNA or ligand binding domains (Figure 1B). This strategy has been shown to be effective at eliminating gene function in larval zebrafish (Hoshijima et al. 2019; Shi et al. 2015). To achieve efficient targeting at specific loci, Alt-R-modified crRNA and tracrRNA were annealed into an RNA duplex and complexed with Cas9 protein (Gagnon et al. 2014; Hoshijima et al. 2019). Assembled dgRNPs were injected into the cytoplasm of one-cell stage embryos. For genes where dgRNPs were designed to target 2 separate sites, 2 dgRNPs were injected in combination. For multigene targeting of gene paralogs or functionally redundant genes, up to 4 dgRNPs (2 dgRNPs for *spi1a/b* and *taz/yap1*; 4 dgRNPS for *bach1a/b* and *junba/b*) were injected in combination.

We first used PCR-based capillary electrophoresis to probe the proportion of indels in dgRNP targeted zebrafish larvae. PCR primers were designed to amplify 100 – 200 base pair (bp) amplicons containing each target site and capillary electrophoresis was used to separate PCR amplicons with 1 – 2 bp resolution (Carrington et al. 2015; Ramlee et al. 2015; Varshney et al. 2015). In 2 day post fertilization (dpf) larvae, the average frequency of non-wild-type sized PCR amplicons (indel frequency) for all target sites was 87.6% (Figure 1C). Out of 28 dgRNPs, *bach1a_2* showed poor activity, achieving an average indel frequency of 17.7%. *junbb_2* (82.3%), *nr1d1_2* (86.7%), *pfkfb1_1* (77.9%), and *rab44_1* (85.9%) dgRNPs resulted in moderate indel frequencies, ranging between 50 and 90%. The 23 remaining dgRNPs achieved high indel frequencies that exceeded 90%. These studies recapitulated previous findings and revealed that all 17 genes were successfully targeted at high-efficiency (>90%) by at least 1 dgRNP in zebrafish larvae.

### dgRNP induced mutagenesis efficiency in adult zebrafish

We next examined the efficiency of dgRNP induced mutagenesis in adult zebrafish. To this end, we raised dgRNP injected crispants to adulthood, and measured indel frequency in adult zebrafish fins by capillary electrophoresis. Fin biopsies have been shown to accurately represent alleles found in somatic tissues and in the zebrafish germline (McKenna et al. 2016; Varshney et al. 2015). Indel frenquency averaged 85.2% at all targeted sites (Figure 1D). Among 28 dgRNPs, *bach1a_2* showed poor activity, 12 dgRNPs showed moderate efficiencies (50-90%), while 15 dgRNPs achieved high efficiencies (>90%). A subset of genes important for adult tissue regeneration are likely required during embryonic development or juvenile growth. We postulated that cells or animals harboring deleterious alleles may be subjected to negative selection, and that wild-type clonal expansion could alter the rates of mutagenesis at adulthood. To test this hypothesis, we compared indel frequencies between larval and adult zebrafish at each of the genomic sites targeted in this study (Figure 1E). For 9 target sites in 7 of the targeted genes, indel frequency was decreased in adults than in larvae (*bach1b, egr1, m17, pfkfb1, spi1b, taz*, and *yap1*). These genes were defined by the indel frequency in one or both of their high-efficiency targeted sites decreasing by >10% from larval to adult stages. In contrast, nr1d1_2 dgRNP was higher indel efficiency in adult animals relative to larvae (Figure 1E). Overall, we found that 82.1% of dgRNPs achieved high indel frequencies in 2 dpf larvae, while 53.6% of dgRNPs maintained high efficiencies in adult fin biopsies (Figures 1F,G). For genes that showed moderate mutagenesis efficiency in adult fish, all targeted crispants were genotyped, and 15 genotyped crispants with the highest indel frequencies were subjected to spinal cord transection injury and subsequent phenotyping (Figure 1H). Due to this selection process, indel frequency averaged 91.1% in animals selected for spinal cord transection. Thus, despite evidence that some mutant alleles are subject to negative selection during animal development and tissue growth, dgRNPs achieve efficient mutagenesis in adult zebrafish crispants.

### Indel frequency by capillary electrophoresis and next generation sequencing

Compared to next generation sequencing (NGS) or restriction enzyme-based genotyping, PCR-based capillary electrophoresis offers a cheaper and less restrictive tool to measure indel frequency (Sentmanat et al. 2018). To confirm the mutagenesis rates derived by capillary electrophoresis, we compared the indel frequencies calculated by capillary electrophoresis to the mutagenesis rates obtained by NGS in individual dgRNP targeted animals at 2 genes. For *dlb_1* and *spi1a_1* target sites, capillary electrophoresis slightly underestimated the prevalence of non-wild-type alleles (Figures 2A and 2B). The discrepancy between indel calculation methods was largely due to small, 1-2 bp indels that were coupled with insertions or deletions of the same size (Figures 2C – 2F). These findings confirmed that capillary electrophoresis measures mutagenesis efficiency with great accuracy, and enables target site selection without the constraint of finding and designing restriction site-based genotyping protocols.

**Figure 2.**
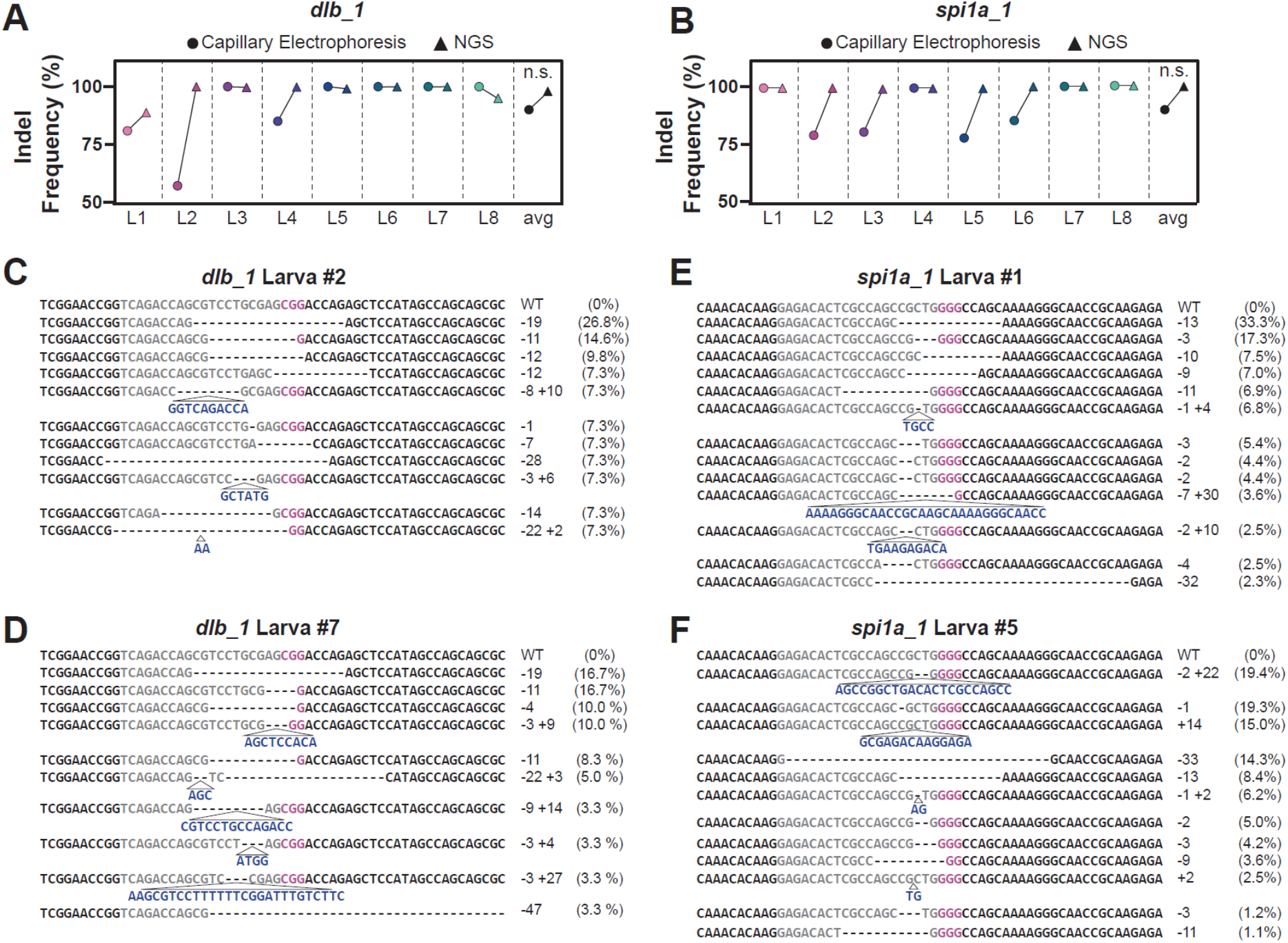
Capillary electrophoresis provides accurate measurement of indel frequencies in zebrafish. **(A,B)** Comparison of indel frequency calculated by capillary electrophoresis (circles) and NGS (triangles) for *dlb_1* (A) and *spi1a_1* (B) target sites in 2 dpf larvae. Eight individual larvae (L1-L8), and the average indel frequency for these larvae (avg) are shown. Statistical analysis was performed; n.s. indicates P-value >0.05. **(C-F)** Sample NGS results for two larvae at *dlb_1* (L2, C and L7, D) and *spi1a_1* (L1, E and L5, F) target sites. Sequence reads represent the wild-type allele (WT) and the mutagenized alleles retrieved at each target site. NGS-derived indel frequencies indicate the percentage of reads mapping to each allele. The target site protospacer is in gray, and the PAM sequence is in magenta. Insertions (navy) and deletions (dashed lines) are indicated.

### Characterization of dgRNP generated alleles

We next investigated the range and diversity of allelic mutations generated by dgRNPs. By NGS, all larvae examined exhibited more than 8 different alleles (Figures 2C – 2E and data not shown). However, only 2-3 alleles were present in >10% of sequenced reads in each animal (data not shown). In 2 dpf larvae, the most common allele was typically present in 20 – 30% of NGS reads for *junbb_1* and *spi1a_1*, and in 10 – 20% for *dlb_1* target site (Figure 3A). These values were consistent with allele prevalence calculated by capillary electrophoresis at all sites (data not shown). Consistent with previous studies, we concluded that dgRNPs were most active at the 2-to 4-cell stage (Burger et al. 2016). In 2 dpf larvae, most alleles were small indels of <10 bp, although larger insertions and deletions were present (Figure 3B). At 3 separate target sites (*dlb_1*, *junbb_1*, and *spi1a_1*), the same indel was generated in multiple, independent animals (Figures 3C – 3E). In *dlb_1* and *spi1a_1* targeted animals, the most common indel shared between independent animals was a frameshift causing deletion (an 11 bp deletion in *dlb_1* [TCCTGCGAGCG] found in 7 out of 8 animals and a 13 bp deletion in *spi1a* [CGCTGGGGCCAGC] found in 6 out of 8 animals, respectively) (Figures 3C and 3D). In *junbb_1* targeted animals, the most common allele was a 3 bp deletion of ACG found in 6 out of 8 independent animals (Figure 3D). Consistent with previous studies, these findings suggested a bias in the double strand break repair process (Ata et al. 2018; Burger et al. 2016; Gagnon et al. 2014), and emphasized the need to target key protein domains to maximize the effect of small indels on gene function.

**Figure 3.**
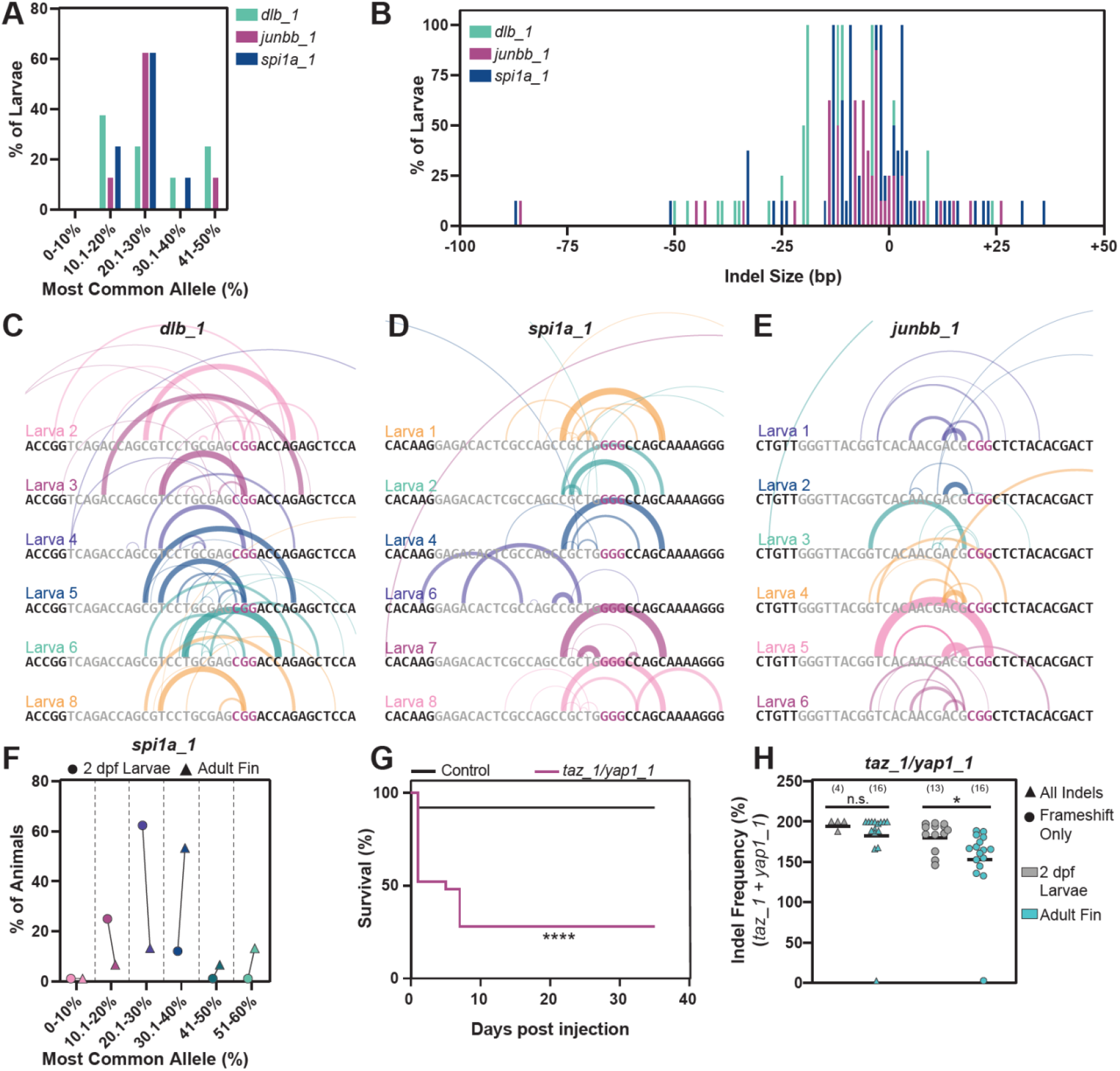
Characterization of dgRNP generated alleles. **(A)** Frequency of occurrence of dgRNP generated alleles for the *dlb_1, junbb_1, spi1a_1* target sites. Data shown represents 8 larvae at each target site and was measured by NGS. Y axis indicates percent larvae; X axis categorizes the most common alleles based on their indel frequency in individual larvae. **(B)** Indel size distribution of dgRNP generated alleles for the *dlb_1, junbb_1, spi1a_1* target sites. Data shown represents 8 larvae at each target site and was measured by NGS. Y axis indicates percent larvae; X axis indicates indel size (ranging from −100 bp deletions to +50 bp insertions) categorizes the most common alleles based on their indel frequency in individual larvae. **(C-E)** Schematics of dgRNP generated deletions in *dlb_1* (C)*, spi1a_1* (D), and *junbb_1* (E) targeted larvae. Individual larvae are shown. Target site protospacers are shown in gray, and PAM sites are shown in red. For each allele, semi-circles indicate base deletion. Line weights and opacity of the semi-circles are proportional to the indel frequency of each allele. The heights of the semi-circle are proportional to deletion size. **(F)** The frequency for the most common allele generated by CRISPR/Cas9 dgRNPs at the *spi1a_1* target site in whole 2 dpf larvae (circles) and adult caudal fin (triangles). Data shown represents eight animals as measured by NGS. The % of animals with its most common allele comprising the particular % of prevalence of reads is shown. **(G)** Survival curves of 25 uninjected wild-type control (black) and 25 *taz/yap1* dgRNP injected (magenta) zebrafish. Sample sizes are indicated. **(H)** The indel frequency of all alleles (triangles) and frameshift only alleles (circles) at *taz_1* and *yap_1* target sites in whole 2 dpf larvae (gray) and adult caudal fins (teal) as measured by capillary electrophoresis. Data points represent individual animals. Indel frequency is shown as the sum of frequencies at the *taz_1* and *yap_1* target sites. Sample sizes are indicated. n.s. indicates P >0.05; (*) P<0.05; (****) P<0.0001

To investigate how allelic prevalence changes from larval to adult stages, we compared indel frequency in *spi1a_1* dgRNP animals at 2 dpf and in adult fin biopsies. While the most common allele in *spi1a_1* crispant larvae was present in 20 – 30% of reads, the most common allele in adults could be found at much higher frequencies (30 – 60%, Figure 3F), suggesting clonal cell expansion occurs throughout development (McKenna et al. 2016). In *spi1a_1* targeted animals, the expanded alleles did not appear to have a bias for in-frame mutations, suggesting clonal expansion may be random for *spi1a*. However, a bias for in-frame mutations was observed for for genes expected to cause early embryonic lethality. Taz and Yap1 are functionally redundant downstream effectors of Hippo signaling (Plouffe et al. 2018). Double homozygous *yap1/taz* zebrafish mutants die during gastrulation (Miesfeld et al. 2015). Due to their functional redundancy, we targeted *yap1* and *taz* alone and in combination. Indeed, 50% of animals injected with both *taz_1* and *yap1_1* dgRNPs arrested prior to 24 hpf and only 30% survived to adulthood (Figure 3G). Among *yap1/taz* targeted adults, significant proportions of wild-type or presumptive in-frame indels (as measured by capillary electrophoresis) were present at one or both *yap1_1/taz_1* target sites (Figure 3H). Taken together, these data support a model in which small indels may minimize the effect of dgRNP induced mutagenesis on certain phenotypes (Gagnon et al. 2014). This may be exacerbated in cases where genes are necessary for development, and mutant cells are removed upon growth to adulthood. Notably, we did not observe this trend in other target sites that were specifically chosen to disrupt key domains. Together, our data suggest that hypomorphic or loss-of-function alleles are considerably prevalent in the majority of genes disruted by in-frame mutations.

### Screening dgRNP targeted crispants for spinal cord regeneration defects

To identify genes that direct spinal cord regeneration, we assessed functional recovery in CRISPR/Cas9 dgRNP injected F_0_ crispants (Mokalled et al. 2016). Mutagenized 3-4 months old animals were subjected to complete spinal cord transection. At 4 weeks post injury (wpi) mutagenized animals and control siblings were subjected to an increasing water current inside an enclosed swim tunnel (Figure 4A). Functional recovery was significantly decreased in *egr1*, *junbb*, *pfkfb1*, *m17*, and *spi1a* crispants relative to their respective control siblings (Figure 4B). Additionally, the defect in functional recovery in F_0_ injected *bach1a/b* crispants was more severe than when *bach1a* or *bach1b* were separately targeted, suggesting *bach1* paralogs are redundantly required for functional spinal cord repair (Figure 4B). Furthermore, *yap1/taz* crispants exhibited a more severe phenotype than *yap1* or *taz* alone (Figure 4B), even though surviving *yap1/taz* adult crispants possessed a bias for in-frame mutations. To rule out that these functional recovery defects were due to developmental or gross morphological defects in targeted crispants, we assessed swim capacity in uninjured *bach1a/b* and *yap1/taz* crispants. In this assay, *bach1a/b* and *yap1/taz* crispants showed comparable swim function to their control siblings, indicating their functional regeneration defects were injury-induced (Figure 4C). These studies revealed dgRNP induced spinal cord regeneration defects in F_0_ crispants.

**Figure 4.**
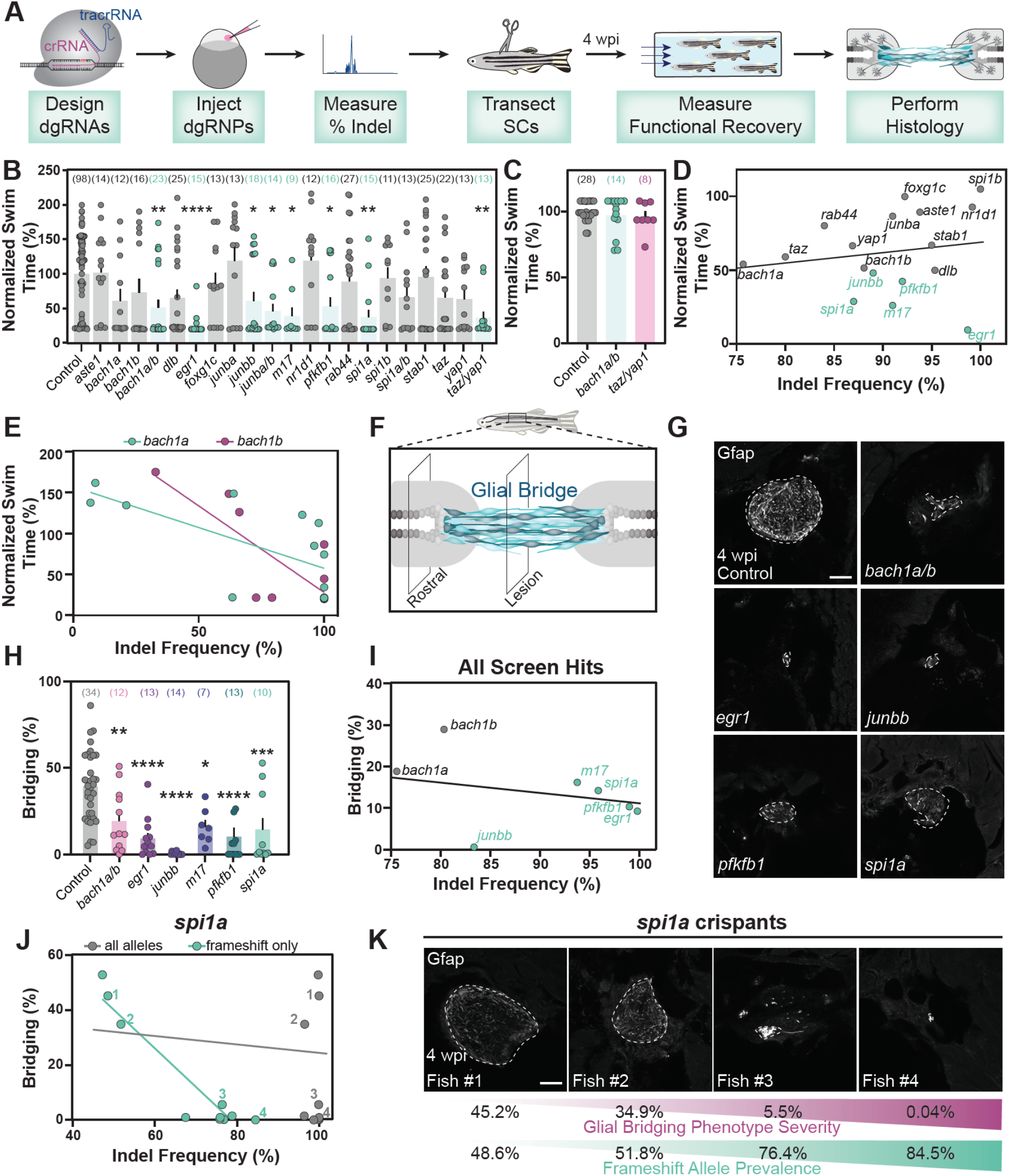
Spinal cord regeneration defects in dgRNP targeted zebrafish. **(A)** Schematic pipeline to screen for spinal cord regeneration phenotypes in dgRNP targeted zebrafish crispants. **(B)** Functional recovery in dgRNP targeted animals 4 wpi. Data points represent individual animals and sample sizes are indicated. For each group of targeted animals, uninjected siblings were subjected to SCI and swim assays. Swim times were normalized to their respective wild-type siblings. Groups with significantly diminished swim function are shown in teal. **(C)** Quantification of swim function in uninjured *bach1a/b* and *taz/yap1* targeted crispants. Data points represent individual animals and sample sizes are indicated. **(D)** Functional recovery plotted against indel frequency (as measured by capillary electrophoresis) for each targeted gene at 4 wpi. Data points represent individual genes. Crispants with statistically significant functional recovery defects are shown in teal. For genes that were targeted at more than one site, indel frequency is averaged across both sites. **(E)** Functional recovery plotted against indel frequency for *bach1a* (teal) and *bach1b* (magenta) dgRNP targeted zebrafish at 4 wpi. Data points represent individual animals. The indel frequency is averaged for target sites #1 and #2, since *bach1a* and *bach1b* were targeted at two sites each. **(F)** Schematic of regenerating zebrafish spinal cord. Bridging glia are shown in teal. Percent bridging was calculated as the ratio of the cross-sectional areas of the glial bridge (lesion) and the area of the intact spinal cord (rostral). **(G,H)** Glial bridging at 4 wpi. Representative immunohistochemistry shows the Gfap^+^ bridge at the lesion site in *bach1a;bach1b*, *egr1*, *junbb*, *pfkfb1* and *spi1a* targeted animals. Percent bridging was quantified for 7-14 animals per group. Data points represent individual animals in H. Sample sizes are indicated. **(I)** Glial Bridging plotted against average indel frequency for each gene (as measured by capillary electrophoresis). Data points represent individual genes. Crispants with significantly diminished bridging are shown in teal. Percent bridging was only measured for crispants that displayed a swim phenotype (B), with the exception of *bach1a* and *bach1b* single crispants. For genes that were targeted at more than one site, indel frequency was averaged across both sites. **(J)** Glial bridging plotted against indel frequency (as measured by NGS) for *spi1a_1* targeted animals. Data points represent individual animals. Gray dots represent indel frequency values of all non-wild-type alleles. Teal dots represent indel frequency of non-wild-type alleles predicted to generate frameshift mutations. Linear regression lines are statistically different (P<0.0001). **(K)** Representative immunohistochemistry shows Gfap^+^ glial bridges in *spi1a* crispants at 4 wpi. Fish 1 through 4 are indicated in panel J. Percent Bridging (magenta) is indicated for each fish along with the frameshift-only indel frequency (teal). For bar plots (B, C and H), the bar indicates the mean ± SEM. (*) P<0.05, (**) P<0.01, (***) P<0.001, (****) P<0.0001. Scale bars, 50 um.

To evaluate the impact of mutagenesis efficiencies on neurobehavioral phenotyping, we compared the extent of functional regeneration relative to indel frequency for different genes and for individual animals per gene. At the gene level, average indel frequency did not correlate with swim capacity after SCI (R^2^ = 0.02) (Figure 4D). We next performed similar comparisons for individual *bach1a* and *bach1b* targeted animals and observed a negative correlation between indel frequency and normalized swim time (R^2^ = 0.45 for *bach1a* and 0.65 for *bach1b*) (Figure 4E). These findings suggest differences in mutagenesis efficiencies among individual animals result in increased variability during neurobehavioral phenotyping.

We next assessed anatomical regeneration in dgRNP targeted crispants after spinal cord transection. After SCI, a pro-regenerative glial bridge is formed and axons regrow across the lesion site (Goldshmit et al. 2012; Mokalled et al. 2016; Becker et al 1997). To measure glial bridging, we imaged and quantified the cross-sectional area of Gfap^+^ bridges at the lesion core relative to the intact spinal cord (Figure 4F). Crispants in *bach1a/b*, *egr1*, *junbb*, *pfkfb1*, *m17*, and *spi1a* displayed less glial bridging (Figures 4G and 4H). To test if indel frequency correlated with glial bridging defects, we compared indel frequencies to percent bridging for different genes and for individual *spi1a* targeted animals. At the gene level, we observed minor negative correlation between indel frequencies glial bridging (R^2^ = 0.14) (Figure 4I), suggesting mutagenesis efficiency may influence phenotypic severity. To specifically explore how frameshift causing alleles may influence phenotypic readout, we performed NGS on spinal cords of individual *spi1a* crispants, and did not observe a correlation between the indel frequency and glial bridging (R^2^ = 0.001) (Figure 4J, gray). However, frameshift-causing indels showed strong negative correlation with percent bridging for frameshift-only mutations (R^2^ = 0.90) (Figure 4J, teal). The 3 *spi1a* crispants with the lowest proportion of frameshift alleles displayed glial bridging comparable with control uninjected siblings (Figures 4J and 4K). The alleles present in the adult spinal cords of *spi1a* targeted animals were remarkably similar to those found in 2 dpf larvae (Figure 3D). The most common *spi1a* allele in adult spinal cords was the same 13 bp deletion present in whole 2 dpf larvae and was found in 10 out of 10 animals present in 5.2 – 29.0% (averaging 19.3%) of sequence reads per animal. However, the second most common allele in *spi1a* crispant adult spinal cords was a 3 bp deletion also found in whole 2 dpf larvae (CTG), present in 9 out of 10 adult spinal cord samples in 2.6 – 29.8% (averaging 10.6%) of NGS reads per animal. The 3 *spi1a* crispant animals with a weaker glial bridging phenotype (Figures 4J and 4K) also had 3 of the 4 highest percentages of reads of the 3 bp CTG deletion. Because the other 7 *spi1a* crispants analyzed possessed a smaller fraction of in-frame alleles and a strong glial bridging phenotype, the weak glial bridging phenotype in the 3 non-frameshift animals did not affect our interpretation. However, we propose genes that cause mild phenotypes may be overlooked due to an occasional lack of frameshift mutagenesis or repetitive generation of non-frameshift alleles. These findings emphasize the importance of using a reliable, sensitive assay to phenotype dgRNP derived F_0_ crispants.

### Generation and neurobehavioral phenotyping of stable germline mutants

To confirm that the phenotypes exhibited by somatic F_0_ crispants were not due to off-target effects, we outcrossed dgRNP injected animals to a wild-type Tubingen background for 2 generations to establish germline mutations. Stable homozygous mutants were then generated for *bach1a*, *bach1b*, *egr1*, *junbb*, and *spi1a* (Figure 5A) (Shaw et al. 2021, In Press). Adult homozygous mutants were subjected to spinal cord transection followed by functional and anatomical phenotyping. At 4 wpi, *bach1a*, *bach1b*, *egr1*, *junbb*, and *spi1a* exhibited functional swim phenotypes with similar severities to their somatic F_0_ crispants (Figures 4B and 5B). Mutants that displayed reduced functional recovery after injury (*egr1*, *junbb*, and *spi1a*) also displayed defective glial bridging at 4 wpi (Figures 5C and 5D). To measure the effect of *egr1, junbb*, and *spi1a* mutagenesis on axon regeneration, we performed anterograde axon tracing and quantified the extent of axon regrowth across the lesion site (Figure 5E)(Mokalled et al. 2016). At 4 wpi, axon regrowth was significantly dimished in *egr1, junbb*, and *spi1a* mutants relative to their wild-type siblings (Figures 5F and 5G). These results confirmed dgRNP targeted F_0_ crispants phenocopy stable germline mutations, suggesting high-efficiency somatic mutagenesis could be used to prescreen for neurobehavioral defects in adult zebrafish.

**Figure 5.**
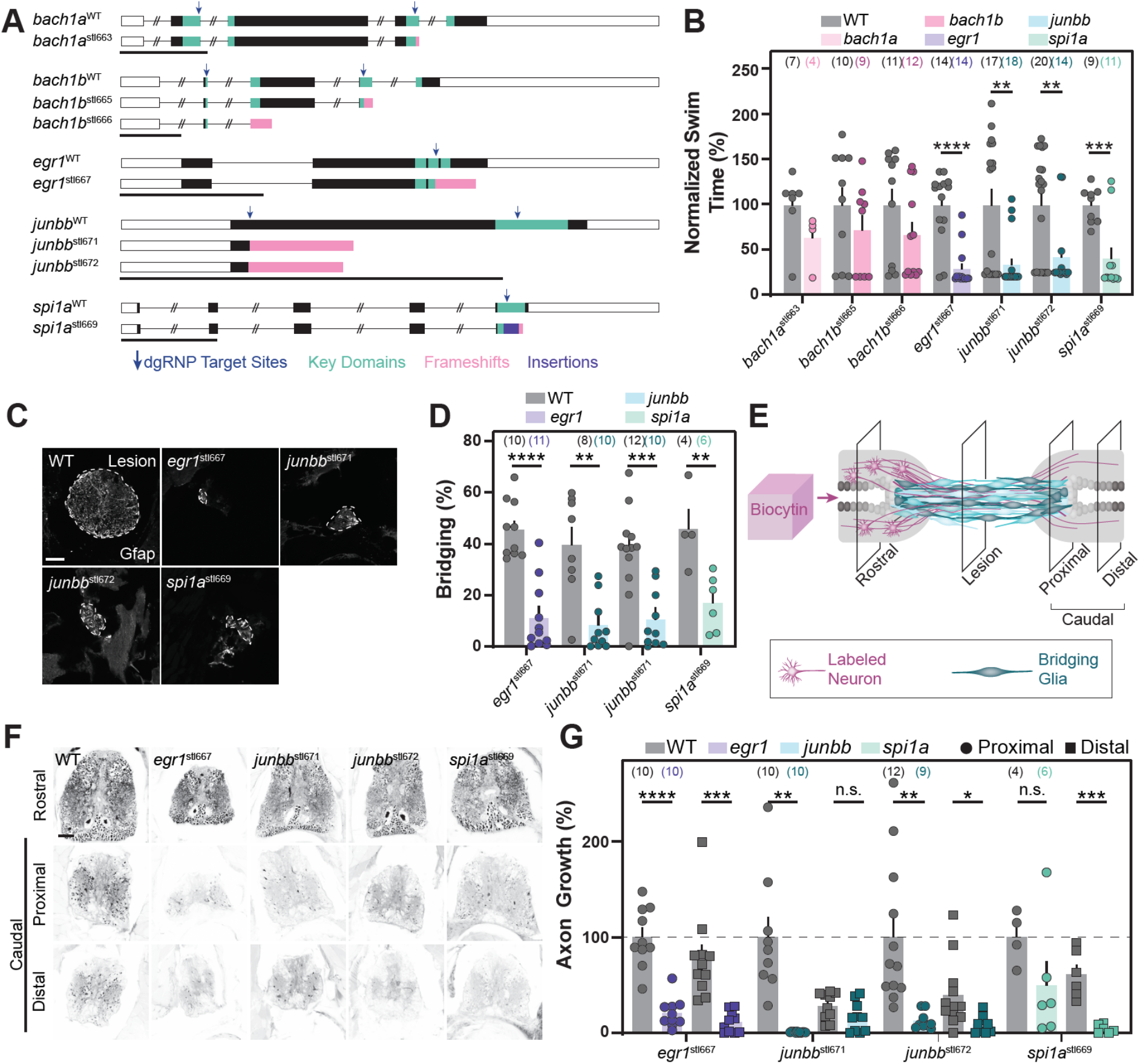
Stable homozygous mutants recapitulate the spinal cord regeneration defects observed in dgRNP targeted crispants. **(A) S**chematics of the germline mutations generated in *bach1a, bach1b, egr1, junbb*, and *spi1a* genes. Black boxes are exons, white boxes are UTRs, and lines are introns. Key domains are indicated in teal. dgRNP target sites are indicated by blue arrows. Predicted frameshift mutations are indicated in pink. A 23 bp insertion in *spi1a* is indicated in purple. **(B)** Functional recovery in stable homozygous mutant lines at 4 wpi. Data point represent individual animals. Sample sizes are indicated. For each clutch of siblings, mutant swim times were normalized to their wild-type siblings. **(C,D)** Glial bridging at 4 wpi. Representative immunohistochemistry shows the Gfap^+^ bridge at the lesion site in *egr1, junbb* and *spi1a* mutants. Percent bridging was quantified for 4-12 animals per group. Data points represent individual animals in H. Sample sizes are indicated. **(E)** Anterograde axon biocytin labeling paradigm. Biocytin labeled neurons (magenta) and bridging glia (teal) are schematized. Axon growth caudal to lesion (proximal and distal) were normalized to the efficiency of Biocytin labeling rostral to the lesion (rostral) for each fish. **(F,G)** Anterograde axon labeling of wild-type control siblings and homozygous mutants proximal (circles) and distal (squares) to the lesion site. Data points in G indicate individual animals. Sample sizes are indicated. For bar plots (B, D, and G), the bar indicates the mean ± SEM. n.s. indicates P >0.05; (*) P<0.05, (**) P<0.01, (***) P<0.001, (****) P<0.0001. Scale bars, 50 um.

## DISCUSSION

This study presents a pipeline to achieve efficient CRISPR/Cas9 mutagenesis and to screen for neurobehavioral defects at medium- or high-throughput in adult zebrafish (Figure 6). To this end, we targeted 17 genes with 28 dgRNPs and achieved somatic mutagenesis efficiencies exceeding 85% in adult animals. Using a quantifiable swim assay as a neurobehavioral readout, we identified 7 genes or gene duplicate pairs that failed to recover swim function after SCI. We show that germline mutations phenocopied somatic crispants, and displayed functional and anatomical defects following injury.

**Figure 6.**
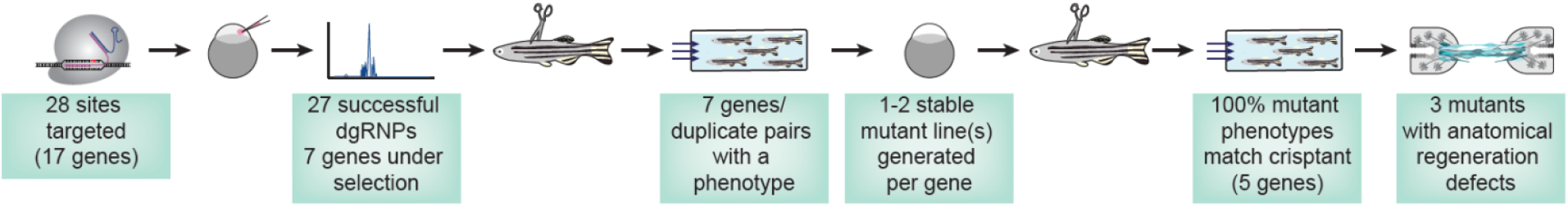
Efficient CRISPR/Cas9 mutagenesis for neurobehavioral screening in adult zebrafish. dgRNPs were used to achieve high-efficiency mutagenesis in targeted crispants. Swim function was used to identify spinal cord regeneration defects after SCI.

To date, performing large-scale genetics in adult zebrafish has proven bulky and challenging. Classical reverse screening techniques rely on easily accessible tissues and highly penetrant phenotypes. We here demonstrate that CRISP/Cas9 dgRNP mutagenesis is highly efficient and reproducible. Thus, parsing through long lists of candidate genes for adult phenotypes is now more feasible. Furthermore, CRISPR/Cas9 dgRNP mutagenesis is cost- and space-effective, making large-scale genetics accessible to even the smallest zebrafish lab. For instance, in our hands, raising 2 tanks of 40 dgRNP targeted adult crispants was sufficient to phenotype each gene in 2 independent experimental replicates. Here, we use a specialized pipeline that our lab has developed for high-throughput screening of spinal cord regeneration factors, but these CRISPR/Cas9-based mutagenesis methods can be combined with a wide array of phenotypic readouts to determine the biological importance of candidate genes in various settings.

Although dgRNPs are an effective tool, there are current limitations to their utility. Small in-frame alleles were generated at all target sites that we sequenced, with differing frequencies. The phenotypic noise generated by these fully or partially functional alleles may cloud the interpretation of certain phenotypes. Consistent with this notion, we found that stable homozygous frameshift mutants in *junbb* displayed a more severe functional swimming recovery phenotype compared to *junbb* crispants (Figures 4B and 5B). The most common allele present in *junbb* crispants was a 3 bp deletion (Figure 3E) that was also recovered in multiple, independent germlines (data not shown). These findings are consistent with this 3 bp deletion being common in adult somatic tissues and likely underlying the difference in phenotypic penetrance between *junbb* crispants and mutants. Targeting essential, functional domains will increase the likelihood of inhibiting or minimizing gene function, but this does not exclude the possibility that mild phenotypes may be missed during screening. Further, the possibility of off-target effects caused by CRISPR/Cas9 dgRNPs cannot be ruled out. Therefore, as we present somatic mutagenesis as an efficient platform to pre-screen for adult phenotypes, confirming the F_0_ crispant phenotypes with multiple, independent germline mutations is indispensable for subsequent phenotypic analysis.

Another limitation to the dgRNP injection methods used here is the inability to achieve temporal or spatial specificity of mutagenesis. To avoid developmental lethality, we pre-screened our candidate list to avoid genes that may cause developmental phenotypes. We took a two-pronged approach to 1) eliminate genes that are known to cause a phenotype incompatible with development to adulthood, and 2) select for genes that are maternally supplied assuming that maternal deposition of mRNA would avoid early developmental defects. Previously, to measure an adult regeneration phenotype, and assuming many genes important for development are also necessary for regeneration, temperature sensitive alleles were generated in a forward genetic screen to dissect fin regeneration (Johnson and Weston 1995; Poss et al. 2002a). Similar temporal or spatial control could be achieved for large-scale reverse genetic screening by optimizing transgenesis of temporally- or tissue-specific reagents. Stable transgenic zebrafish lines driving the expression of Cas nucleases under the control of tissue- or temporal-specific promoters and sgRNAs under the control of the *U6* promoter could attain finer control of Cas-dependent mutagenesis. A recently published study used a tissue-specific promoter to drive the expression of Cas9 in zebrafish astrocytes and achieved tissue-specific gene knockout of a target gene (Chen et al. 2020). However, more work on these systems is required to achieve indel generation rates high enough to measure quantitative and variable readouts, such as swim function after SCI.

Adult zebrafish are a leading vertebrate system to model human diseases and dissect tissue regeneration mechanisms. However, the anatomy of the regenerating spinal cord in adult fish is complex and requires prolonged histological processing to acquire and analyze tissue architecture. By using a functional swimming assay to pre-screen fish prior to histology, we were able to successfully identify 7 novel genes or gene-duplicate pairs that direct functional spinal cord repair after injury. A three-pronged approach that combines transcriptomics, reliable CRISPR/Cas9 dgRNP mutagenesis, and robust functional phenotyping, proved to be a powerful approach toward neurobehavioral phenotyping in adult zebrafish.

## METHODS

### Zebrafish

Adult zebrafish of the Tubingen strain were maintained at the Washington University Zebrafish Core Facility. All experiments were performed in compliance with institutional animal protocols. Male and female animals between 3 and 6 months of ~2 cm length were used. Experimental fish and control clutch-mate siblings of similar size and equal sex distribution were used for all experiments. Spinal cord transection surgeries and regeneration analyses were completed in a blinded manner, and 2 independent experiments were repeated using different clutches of animals.

### CRISPR/Cas9 Mutagenesis

CRISPR/Cas9 design and mutagenesis was performed as previously described (Hoshijima et al. 2019). Briefly, crRNA guide RNA sequences were selected using CHOPCHOP (https://chopchop.cbu.uib.no/). Only sequences with no off-target sites with three or fewer mismatches elsewhere in the genome were selected. To maximize the effect of small indels, target sites were chosen that lie within essential domains. For genes that were targeted twice, an additional second target site was selected within an early exon. Target sequences used for this study are outlined in Supplementary Table 1.

Alt-R tracrRNA and crRNA gRNAs (IDT, Cat# 1072534) arrived as lyophilized powder and were reconstituted using manufacturer’s specifications as 100 uM stocks and stored at −20°C. Prior to the day of injection, crRNA and tracrRNA were mixed at a final concentration of 50 uM and annealed by heating to 95°C and then gradual cooling to 25°C (−0.1°C/second). The resulting dgRNA duplexes were stored at −20°C until use. Alt-R S.p. Cas9 nuclease V3 (IDT, Cat# 1081059, supplied at 61.7 uM in 50% glycerol) was diluted by Cas9 dilution buffer (1 M HEPES (pH 7.5), 2 M KCl) to a working concentration of 25 uM and stored in single use aliquots at −80°C. On the day of injection, annealed dgRNA duplexes were diluted 1:1 in duplex buffer (IDT, Cat# 11-05-01-03) to a working concentration of 25 uM. Equal volumes of dgRNA were added to Cas9 protein and incubated at 37°C for 5 minutes. For samples where 2 (*aste1, bach1a, bach1b, dlb, junba, junbb, nr1d1, m17, pfkfb1, rab44, spi1a/spi1b, stab1*, and *taz/yap1*) or 4 (*bach1a/bach1b* and *junba/junbb*) sites were being targeted at once, Cas9 protein was added in equal molar amounts to the total concentration of dgRNA. CRISPR/Cas9 solutions were maintained at room temperature during injections. Tubingen wildtype embryos were injected with 1 nL of CRISPR/Cas9 solution at the one-cell stage and grown to adulthood for spinal cord surgeries and functional analysis.

### Capillary Electrophoresis

Capillary electrophoresis (fragment analysis) was used to calculate the indel frequency for each CRISPR/Cas9 target site. For DNA extraction, whole 2 dpf larvae or ~3 mm of excised adult tail fins was added to 50 mM NaOH in 50 uL (larvae) or 100 uL (adult fin). DNA samples were incubated at 95°C for 20 minutes and then rapidly cooled to 4°C. DNA extractions were neutralized by adding 5 uL (larvae) or 10 uL (adult fins) of 1 M Tris-HCl (pH 8.0). Small 100-200 bp PCR products were generated using NEB Taq Polymerase (Cat# M0273) with gene-specific primers (Supplementary Table 2) in a volume of 10 uL in Pryme PCR semi-skirted PCR plates (MidSci, Cat# AVRT1). Samples were diluted to 24 uL with TE dilution buffer (Agilent) and loaded into the 5200 Fragment Analyzer System (Agilent, Cat# M5310AA). Capillary electrophoresis was carried out using the Agilent Fragment Analyzer Qualitative DNA Kit (Cat# DNF-905-K1000) according to the manufacturer’s specifications.

To calculate indel frequency, three wildtype uninjected siblings were also PCR amplified. The size of wild-type products was determined using these control samples. Any peaks within 1 bp of the wild-type control amplicons was called as wild-type (non-indel). The indel frequency was calculated by dividing the total non-indel peak signal by the total peak signal. Because of significant noise due to primers (<70 bp) and non-specific products (>200 bp) in the wild-type samples, only signal between 70 and 200 bp was used to calculate indel frequency. Capillary electrophoresis primers were also used for genotyping stable mutant lines.

### Next Generation Sequencing

DNA extracts from larvae or adult fins were directly submitted to the Washington University Genome Engineering & iPSC Center (GEiC) for next generation sequencing on the 2×250 Illumina MiSeq platform (Sentmanat et al. 2018). The same primer sets as for capillary electrophoresis were used for NGS.

### Surgeries

Adult fish were anesthetized in 0.02% tricaine and placed in a shallow slit cut into a damp cellulose sponge, dorsal side facing up. Fine tip forceps were used to remove scales 3 mm rostral to the dorsal fin. Microscissors were used to cut (perpendicular to the rostral/caudal axis) through the overlying skin and muscle. A second cut was used to fully transect the spinal cord. Forceps were used to check the injury site and ensure the spinal cord was completely transected. Injured fish were placed in clean system water to recover and a transfer pipette was used to flush water through the gills to aid in recovery from tricaine anesthetic.

### Immunohistochemistry

Adult fish were euthanized in 0.02% of tricaine and their spinal cords dissected and fixed overnight at 4°C in 4% paraformaldehyde (PFA) at indicated timepoints. Spinal cords were washed out of PFA into PBT (0.1% Tween-20 in PBS) and incubated with rocking in 30% sucrose overnight at 4°C. Spinal cords were embedded in OCT freezing media and frozen blocks were stored at −80°C until sectioning into 25 um transverse cryosections. For Gfap staining, sections were rehydrated into PBT and then incubated overnight at 4°C in mouse anti-Gfap (ZIRC, Zrf1) diluted 1:1000 in blocking agent (5% goat serum in PBT). Sections were washed in PBT and incubated for 2 hours at room temperature in Alexa Fluor-488 anti-mouse secondary antibody (Invitrogen) diluted 1:250 in blocking agent. Sections were washed in PBT and mounted in Fluoromount-G media.

Anterograde axon labeling was performed on adult fish at 4 wpi. Fish were anaesthetized in 0.02% tricaine and fine scissors were used to transect the spinal cord 4 mm rostral to the lesion site. 1 mm^3^ of Gelfoam gelatin sponge (Pfizer, Cat# S-11227) was soaked with 1.25 uL of 40 mg/mL biocytin (Sigma, Cat# B4261) and inserted into the new lesion site. Vetbond tissue adhesive (3M) was used to close the incision. Fish were euthanized 4 hours post-labeling by tricaine overdose and fixed, dissected, and sectioned as described above. Sectioned spinal cords were rehydrated into PBT, stained for 2 hours at room temperature with Alexa Fluor-594-conjugated streptavidin diluted 1:100 in 0.1% triton X-100 in PBS, washed in PBS, and mounted in Fluoromount-G media.

### Quantification and Statistical Analysis

All procedures and quantifications were performed blind to condition. To calculate functional swimming recovery (Normalized swim time; Figures 4B, 4C and 5B), the swim time for each individual fish was divided by the average swim time for its wild-type or uninjected control clutch-mate siblings and multiplied by 100. To calculate glial bridging (Bridging %); Figures, 3H and 4D), the cross-sectional Gfap^+^ area of the center of the lesion was divided by the cross-sectional area of the spinal cord 1 mm rostral to the lesion site and multiplied by 100. To calculate the Axon Growth (Figure 3G), biocytin labeled axons were quantified using the threshold and particle analysis tool in Fiji. Multiple sections for each fish at each level were quantified: +0.5 mm (caudal – proximal, 4 sections), +2 mm (caudal – distal, 4 sections), −1 mm (rostral, 2 sections). The number of biocytin^+^ particles in each region of the spinal cord (caudal – proximal, caudal – distal, and rostral) were averaged for each individual fish. The caudal – proximal and caudal – distal values were divided by the rostral value to give the axon growth index. This value was then normalized to the average caudal – proximal value in wild-type sibling controls and multiplied by 100.

Statistics were performed in GraphPad Prism. In cases where two groups were compared (Figures 3H, 5B, 5D and 5G), t-tests with Welch’s correction (where appropriate) were used. When three or more groups were compared (Figure 4B, 4C and 4H), One-way ANOVA with Dunnett’s correction was used. For the survival analysis (Figure 3G), a Log Rank Mantel-Cox test was used. To compare linear regressions in Figure 4J, an Anova test was used.

---

**Supplementary Table 1.**
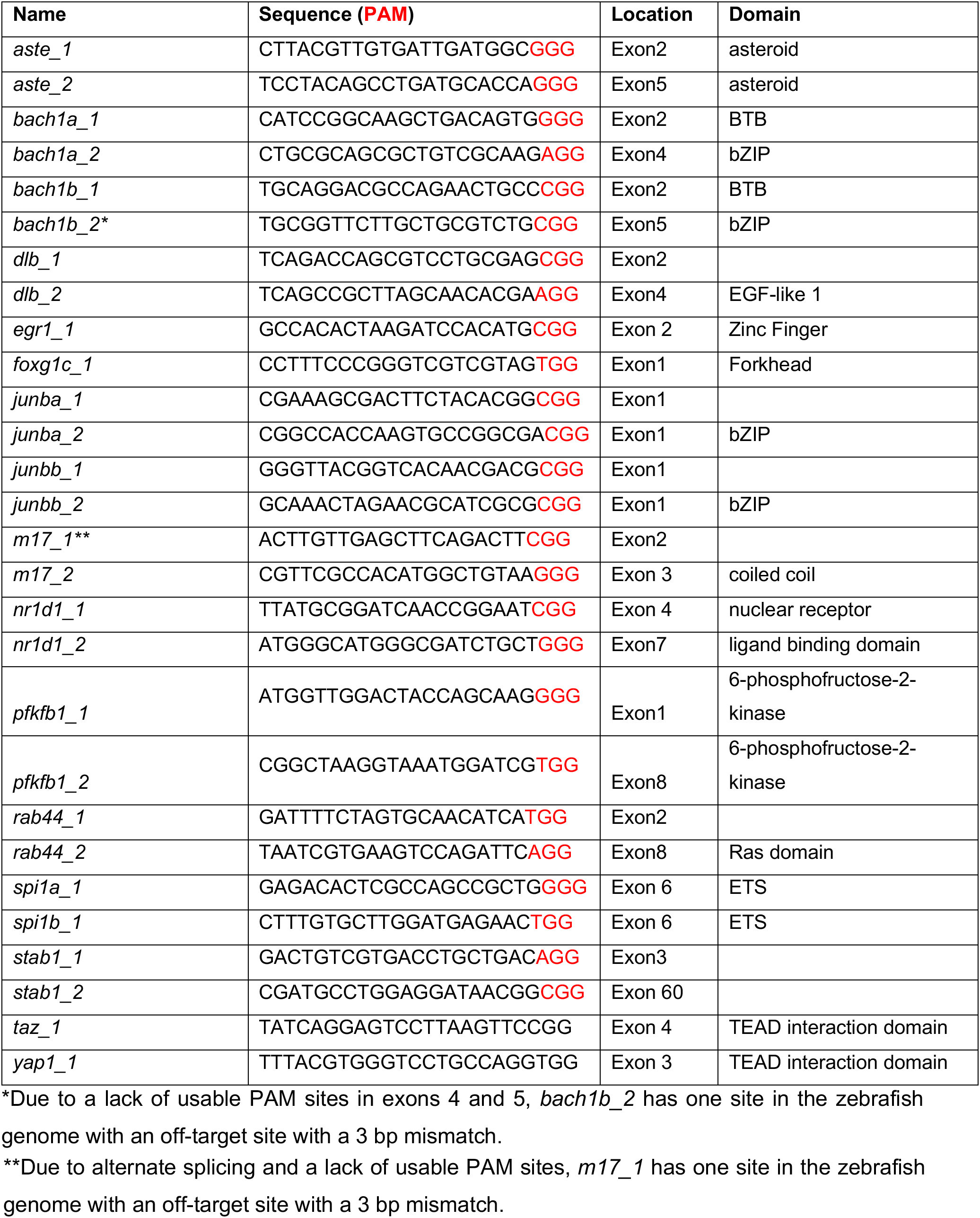
CRISPR/Cas9 target sites

**Supplementary Table 2.**
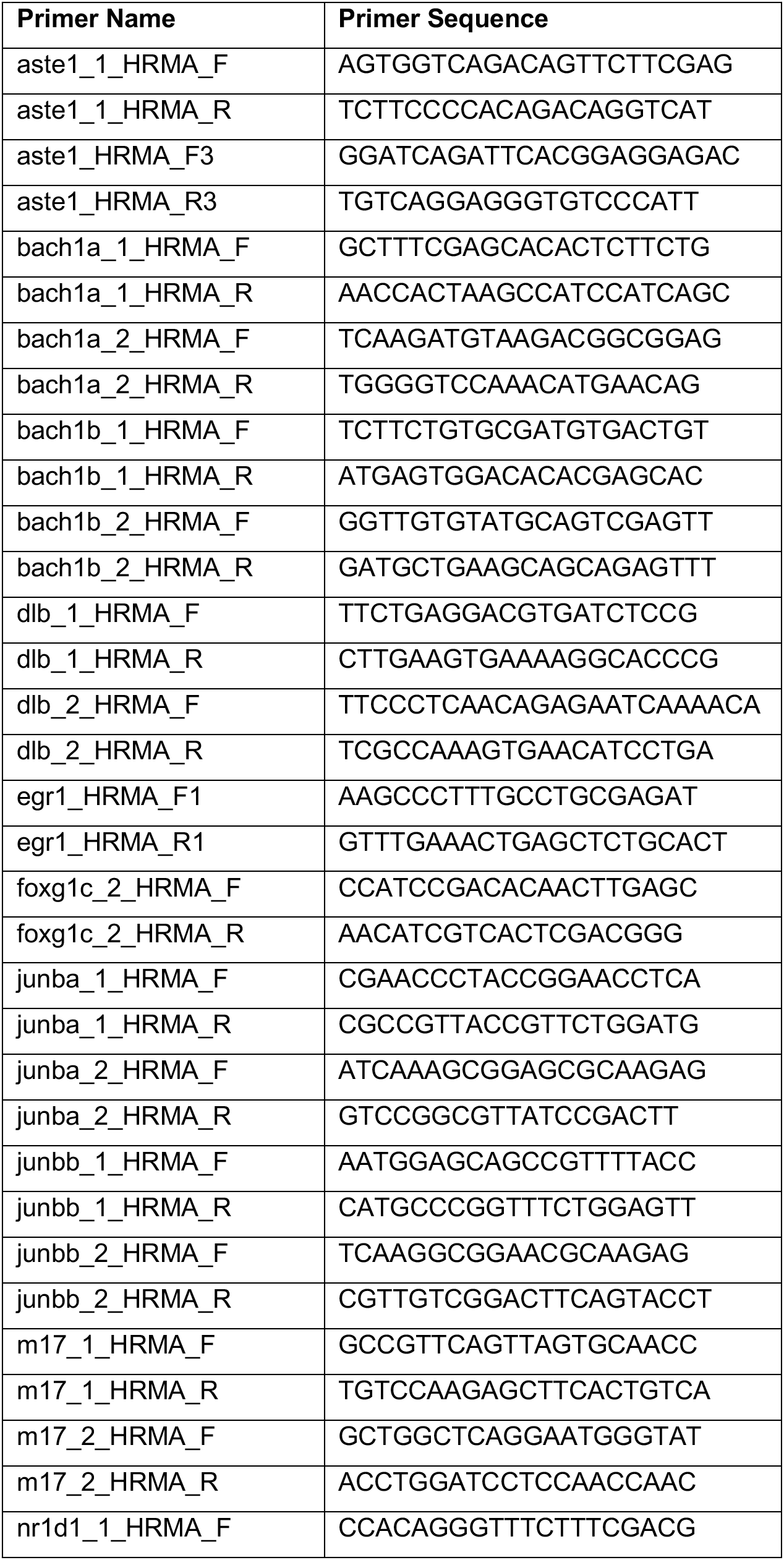
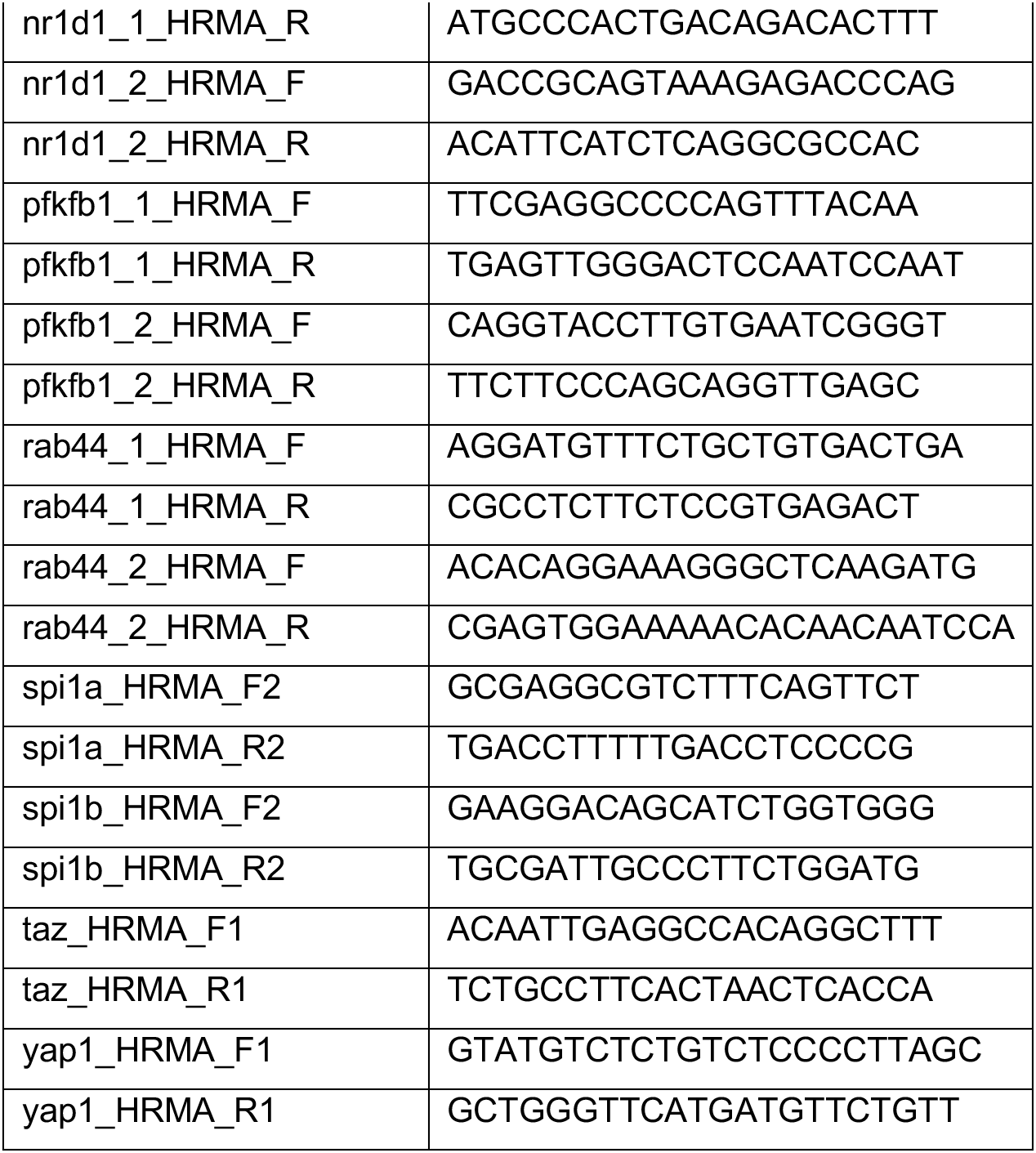
Genotyping primers used in this study

